# Combining Multi-site FRAP and HILO-TIRF microscopy using a Spatial Light Modulator

**DOI:** 10.1101/2025.04.15.649016

**Authors:** Avinash Upadhya, Yean Jin Lim, Woei Ming Lee

**Author notes:** &.

## Abstract

Fluorescence Recovery After Photobleaching (FRAP) has remained a powerful tool to probe intracellular dynamics. FRAP relies on two aspects: (1) localised excitation resulting in photobleaching, and (2) fluorescence recovery of the bleached volume which provides insight into kinetics. Existing FRAP systems are limited by a trade-off between time and spatial multiplexing due to galvanometric scanning methods, precluding the study of multiple independent positions simultaneously as well as advanced widefield imaging modes. Hence, they may not capture the dynamic and non-isotropic environments in biological studies. In this paper, we utilise phase profiles corresponding to an array of fresnel lenses and diffractive masks to switch between simultaneous FRAP of independent spatial positions whilst maintaining epifluorescence, highly inclined laminated optical sheet (HILO), and total internal reflection fluorescence (TIRF) microscopy modalities. Our approach bridges high contrast fluorescence imaging (HILO and TIRF) and a multi-position FRAP technique using a single spatial light modulator. As such, this technique enables high contrast bleaching and screening across volumes, which we envision will be of value to areas such as single particle tracking and single molecule imaging where dynamic photobleaching is necessary to measure fast events across the field of view in a single versatile instrument.

## 1. Introduction

From its conception, the development of FRAP Fluorescence Recovery After Photobleaching (FRAP) has been strongly influenced by the signal to noise ratio of the fluorescence imaging instrumentation [1]. FRAP has been used since the early 1970s in custom built optical instruments as well as in Confocal Laser Scanning Microscopes (CLSMs) [1–4]. The early studies by Liebman and Entine [5], and Poo and Cone [4] illustrated the concept of photobleaching and recovery in a biological sample. These studies established that not only were such proteins freely diffusing in the membrane, but that they could be irreversibly photobleached. However, Peters et al [3] showed that recovery does not always follow bleaching, which they attributed to restricted diffusion due to protein aggregation and a ‘rigid’ membrane. Recently, FRAP has been utilized to determine the diffusion rates and viscosity of condensates within membrane-less intracellular structures for phase separation biology [6]. Such applications require live imaging at sub-cellular resolutions that exceed the capabilities of confocal microscopy.

Highly inclined laminated optical sheet (HILO) is a technique that operates in between Total Internal Reflection Fluorescence (TIRF) and epifluorescence microscopy [7]. HILO is compatible with live, sub-cellular imaging, yielding an SNR eight times greater than epifluorescence, while being suitable for single molecule and single particle tracking microscopy [7]. TIRF, however, illuminates only regions adjacent to the coverslip, and has the best axial resolution of the techniques [8, 9]. These methodologies make use of the same optical setup, with a key difference being the position of the focus in the BFP and hence angle of illumination at the sample. However, standard HILO microscopes are not inherently designed for the precise, localized photobleaching required for many FRAP experiments [10]. This makes seamless switching between HILO/TIRF imaging and localized FRAP difficult. Introduction of additional beams can alleviate this issue albeit at the cost of increased system construction and alignment complexity. Furthermore, faster alternatives to galvanometric mirrors such as acousto-optic modulators (AOMs) and acousto-optical tunable filters (AOTFs) are a possible solution, but can be limited by diffraction efficiency and timing constraints and do not permit axial control.

In this paper, we sought a versatile solution to combine the need for FRAP incorporating both spatial and temporal information with high contrast imaging methods like HILO and TIRF. Here, we present a system that unifies inclined imaging modes with multi-spot FRAP through the use of a nematic Liquid Crystal on Silicon (LCoS) spatial light modulator (SLM) [11]. SLMs act as programmable diffractive optical elements within the optical train, and have been used widely in optical microscopy [12].

## 2. SLM enables real-time switching between widefield inclined imaging and multi-spot photobleaching

CLSM reinvigorated the FRAP technique, particularly in application to live cell protein studies [1, 13, 14]. FRAP has now been conducted in virtually every compartment within the cell [1]. While there are other techniques that can be used to study diffusion dynamics such as Fluorescence Correlation Spectroscopy (FCS) and Single Particle Tracking (SPT), these can require non-standard optical instrumentation and sample preparation [15]. The flexibility and wide availability of galvanometric mirrors in CLSMs was key to the proliferation of FRAP, where angular rotation of the mirrors steers a beam focus and photobleaches fluorescent regions as in Figure 1 A). This optical configuration, however, has several drawbacks.

**Fig. 1.**
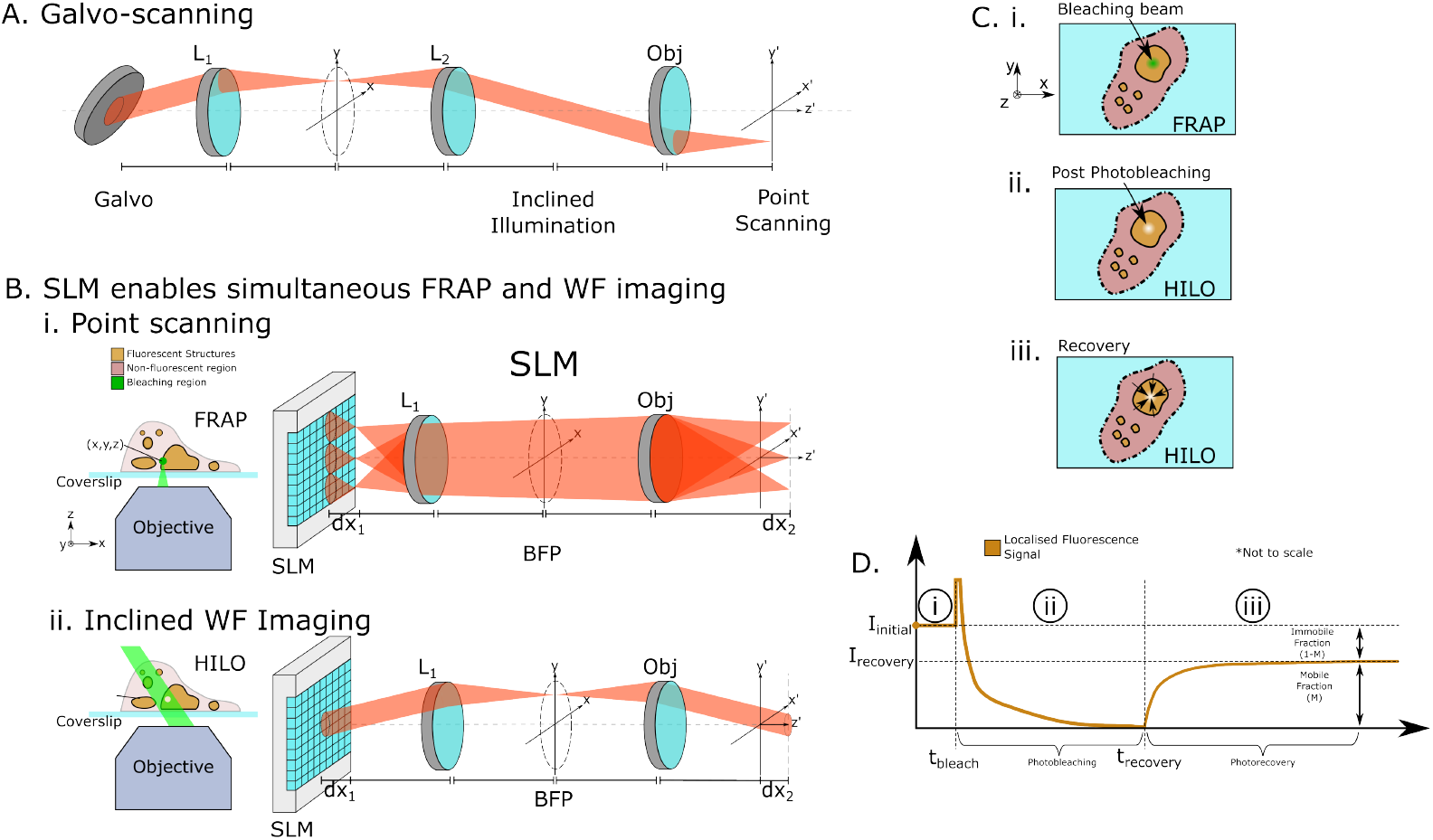
A point scanning FRAP system using scanning mirrors. A) The angular positions of the galvanometric mirror determines the bleaching position, precluding use of inclined illumination at the same plane. B) As a versatile diffractive optical element, the SLM enables simultaneous i) FRAP at multiple points and ii) widefield imaging. This allows interrogation of biological samples using both FRAP and inclined widefield imaging modes such as HILO in the same optical train. C) illustrates the use of such a system for FRAP, with i) bleaching of a cellular sample in a FRAP system, ii) resulting photobleached region observed by HILO, iii) subsequent recovery by observing influx of fluorescent molecules into the compartment. The fluorescence recovery curves at this point us shown in D). In D) i) the initial fluorescent intensity I_initial_ is measured using a lower power imaging beam showing constant intensity. In D) ii) at t_bleach_, the high power bleaching beam is switched on leading to a momentary increase in fluorescent intensity (not to scale) followed by decrease as molecules are photobleached. In D) iii) bleaching beam is switched off and imaging beam is switched on again leading to increase up to I_recovery_. I_recovery_ < I_initial_ as a subset of fluorescence molecules are attached to objects of interest that are bound in the region.

Firstly, the spatial profile of the bleaching beam itself can influence mathematical models of diffusion and kinetics. Some studies have modified CSLMs to bleach regions of different shapes such as disks and lines [16, 17], or in combination with computational techniques [18]. The Gaussian point scanning geometry of most CSLMs restricts this, where the optical train in Figure 1 A restricts illumination at the objective focus to a single point focus. This precludes the spatial multiplexing of bleaching spots. This has implications for application of FRAP to complex biological samples where diffusion constants can vary across the field of view (FOV). While time sharing can be employed to generate a number of bleaching spots, this can become limited by even high speed scanning mirrors. Moreover, galvanometric scanning mirrors cannot achieve axial control over the foci which would require lens elements. Similarly, the illumination optics in HILO (and TIRF) microscopes rely on widefield inclined illumination at the sample and are not inherently designed for the localized photobleaching required for FRAP experiments. This makes it difficult to switch between the these two modalities.

We implemented an SLM to bridge the optics between inclined imaging modes and FRAP. Figure 1 B shows the use of a nematic SLM in our optical train to tailor excitation. In Figure 1 B) i) the SLM permits individual, simultaneous control of multiple foci at the sample plane, which is used for bleaching in FRAP. Characterisation of the bleaching spot position formation are presented in Supplementary Figure S1. In Figure 1 B) the same optical train and SLM is used to incline the excitation beam at the sample in application to HILO and TIRF imaging. Hence the SLM can allow FRAP measurements at different points in a cell simultaneously, while observing the sample through HILO or TIRF imaging. Switching between the two modalities is instantaneous, allowing observation of cell motion across the FOV, which will be important for correlative studies in cell biology.

Figure 1 C) describes the FRAP process at a three bleaching spots modulated by the SLM. In Figure 1 C) i), a focused beam (green) bleaches the fluorescence within an organelle (orange) with the SLM in FRAP mode. Subsequent bleaching can be observed by HILO imaging in the lighter region of Figure 1 C) ii). Finally recovery of fluorescence intensity as bleached molecules are exchanged with unbleached molecules as in Figure 1 C) iii). Figure 1 D illustrates the fluorescence intensity in FRAP in regions corresponding to Figure 1 C. HILO is first used to measure baseline fluorescent intensity I_intensity_ in a localised region. At t_bleach_, a high power bleaching beam causes a momentary increase in fluorescence, followed by exponential decrease in fluorescence signal. At t_recovery_, HILO is switched on once again in region C, causing an exponential recovery of fluorescence as bleached molecules are replaced by unbleached molecules. This results in final steady state intensity I_recovery_ being reached asymptotically with time. It is noted that I_recovery_ is typically lower than I_initial_. If we consider that the I_initial_ denotes the number of fluorescent molecules, we can break this down into the mobile fraction (M) and immobile fraction (1-M). The mobile fraction represents those fluorescence molecules which are unbound and freely diffusing. The immobile fraction indicates a portion of bleached molecules are bound in the region, and so cannot be replaced.

We describe the types of back focal plane (BFP) phase profile necessary to enable switching between the imaging and FRAP modes in Figure 2. Figure 2 illustrates that switching between epifluorescence, HILO, and TIRF methods is based on restricting light to radial regions in the objective BFP. As such in Figure 2 A) the BFP of a high NA objective is segmented into regions, epifluorescence corresponds to a centred beam spot (grey region), HILO to off-axis positions (yellow region), and TIRF to the edge of the objective (red region). While switching between epifluorescence, HILO, and TIRF is usually achieved through scanning mirrors, this is not possible in typical point scanning optical trains as in Figure 1 A). We changed the phase mask displayed on the SLM to scan the spot in the BFP and hence emerging inclined excitation (shown in green), switching between epifluorescence, HILO, and TIRF modes as shown in Figure 2.

**Fig. 2.**
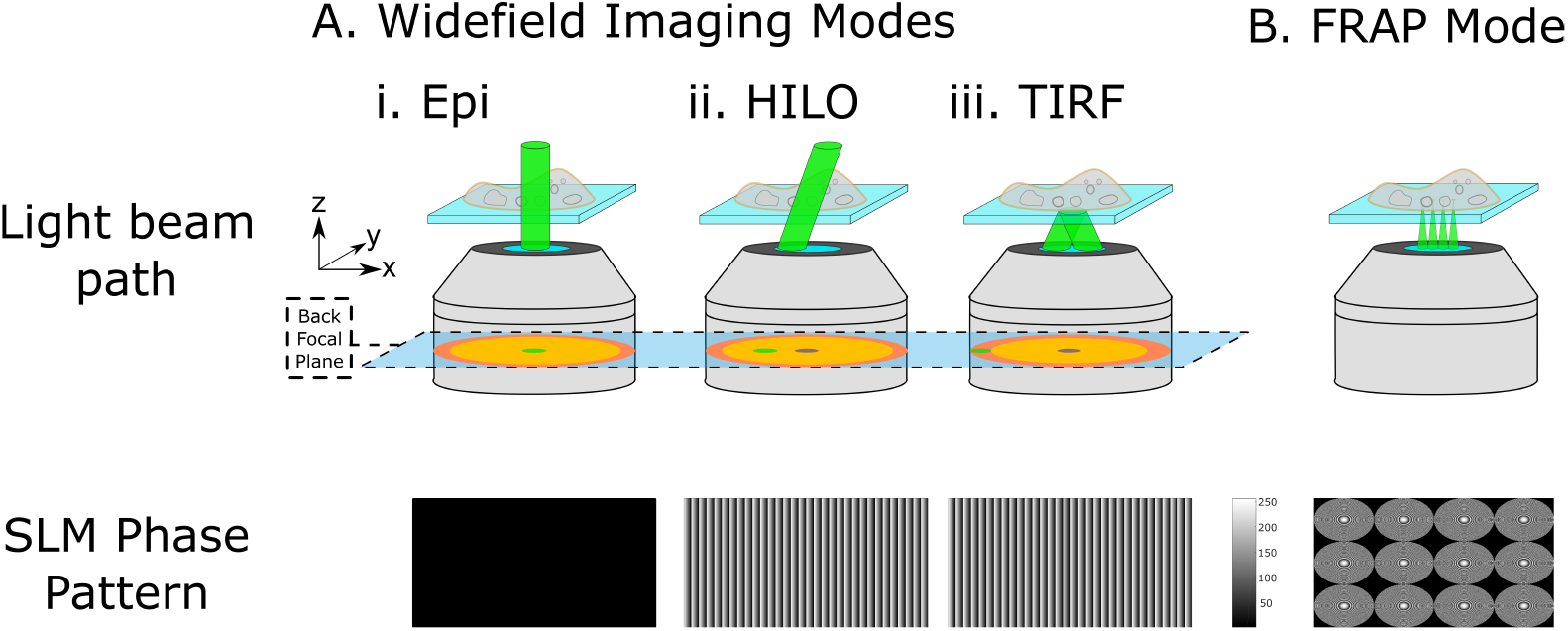
Illustration of widefield fluorescence modalities using high NA objective A) i) epifluorescence excites regions of a sample along the axial direction (BFP spot oriented to optical axis), ii) HILO inclines the beam to reduce out of focus excitation and higher SNR (BFP spot translated). iii) close to the edge of the BFP, a highly inclined beam with *θ* > *θ*_*critical*_ leads to total internal reflection. The evanescent wave excites only fluorophores close to the coverslip (<150 nm). This gives the highest SNR but cannot image deeper structures. B) Usage of a micro lens array phase pattern on the SLM generates numerous steerable foci at the sample plane.

In Figure 2, we illustrate the blazed grating phase profiles *ϕ*_*blazed*_ = *T xcos* (*θ*) displayed on the SLM to achieve HILO and TIRF, where *T* is the blaze parameter controlling radial position of the spot, and *x* is an SLM coordinate, and *θ* is the ramp direction relative to the x-axis. Figure 2 A) i) shows a uniform SLM phase corresponding to epifluorescence. Figure 2 A) ii) and Figure 2 A) iii) show two blazed gratings used for HILO and TIRF respectively at T=-600 and T=-680.

In addition, multiple foci can be generated at the objective focus using SLM with phase mask consisting of an array of Fresnel micro lenses as in Figure 2 B). Each lens follows phase with form 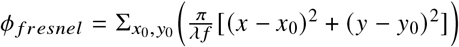, where (*x, y*) are coordinates in the SLM plane, (*x*_0_, *y*_0_) are microlens positions, *λ* is excitation wavelength, and *f* controls micro lens focal length. Hence, segmented regions of pixels correspond to the individually addressable foci (see Supplementary Figure S2). Each microlens can be tuned through focal length or with additional phase, as well as switched off by replacing its SLM segment with a flat phase profile. In Figure 2 B), a phase mask capable of projecting 3× 4 foci is shown along with the resulting microscope image. Foci are formed close to the focus of the objective (~ *μ*m away), with small movement of the objective and camera position used to collocate the foci at the objective BFP.

Our system used a high NA objective lens to enable a greater range of beam inclinations. TIRF requires beam inclinations *θ* > *θ*_*c*_ = *sin*^−1^ (*n*_2_ /*n*_1_) where *n*_1_ is the refractive index of the coverslip and *n*_2_ is the refractive index of the sample (typically water) [19]. This condition sets a minimum objective NA of 1.40 but NA>1.45 is typically utilised [20]. Higher NA objectives above this point permit a greater range of angles above the critical angle, thereby allowing modulation of the penetration depth of evanescent excitation. For example, objectives are now available with NA up to 1.7 which can be used for TIRF [21].

Figure 3 A) illustrates mode switching functionality on fixed L929 mouse fibroblast cells stained for F-actin using phalloidin and excited by 488 nm light, with an Olympus NA 1.49 objective. In Figure 3 A) i) epifluorescence image shows out of focus fluorescence intensity. In Figure 3 A) ii), the HILO image still shows structures deeper within the cell, but with a higher SNR (the haze is reduced). In Figure 3 A) iii), the TIRF image excites only those regions adjacent to the coverslip. In this case no deeper objects are excited leading to the highest SNR. We switch modalities using the SLM via a custom written software interface (see Supplementary Figure S3).

**Fig. 3.**
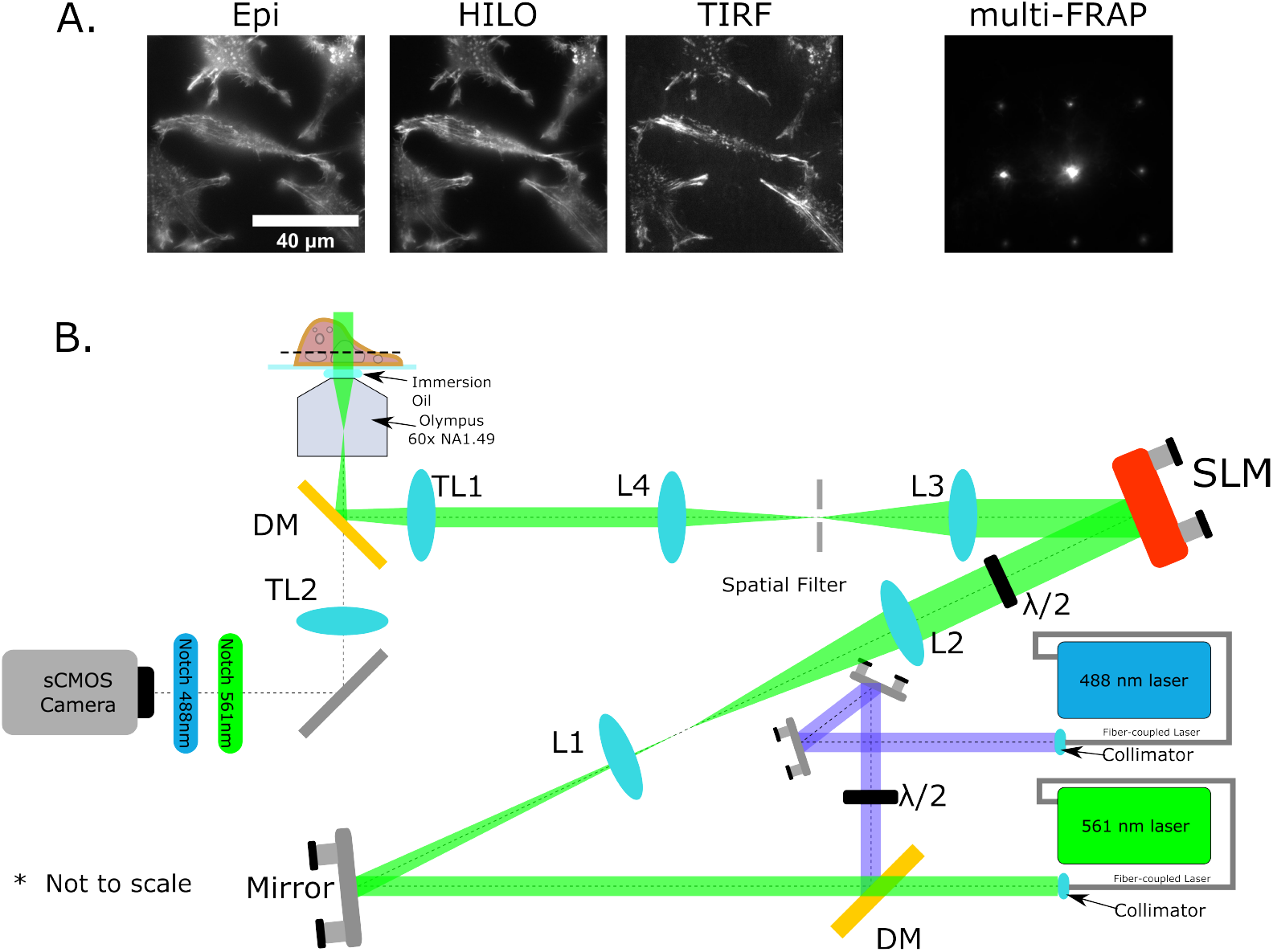
A) Fixed L929 mouse fibroblast cells stained for F-actin using phalloidin (488 nm excitation) in epifluorescence, HILO, TIRF, and multifocal excitation. TIRF has the highest Signal to Noise, while HILO maintains the ability to image structures further from coverslip. B) Optical schematic for HILO-FRAP system. Two fibre coupled lasers (488 and 561 nm) lasers are expanded, filtered (as necessary), and coaligned using a dichroic mirror DM. Once coaligned, both beams are directed by a mirror towards a pair of expansion lenses L1 and L2 (we omit the 488 nm beam for clarity). SLM reflects beam towards L3, which focuses at the Fourier plane, where spatial filtering is conducted via iris and zero order block. Light is then recollimated by L4, and directed towards TL1, which reforms the Fourier plane in the BFP of the objective lens through the dichroic mirror DM, leading to collimated beam at sample. Fluorescence emission of stained sample is transmitted by the dichroic mirror, and TL2 images the sample plane at the camera.

The final optical design is shown in Figure 3 B). The system comprises two fiber coupled semiconductor lasers of 561 nm (Cobolt) and 488 nm (Coherent). Each laser was collimated, spatially filtered, and expanded to give a well collimated beam that maintained beam diameter over at least six metres of propagation. A dichoic mirror (Semrock) coaligned the beams, with subsequent beam expansion to overfill the SLM and half waveplate to ensure linear polarisation parallel to SLM extraordinary axis. Optimal linear polarisation is critical to achieving high first order efficiency. The SLM then reflects excitation light at small angle (< 10 degrees) towards L3, and spatial filtering was conducted at its focal plane using an iris to block higher orders, and a static amplitude filter (adhesive putty on coverslip) blocking the zero order light. Positioning of spatial filtering was conducted using three-axis stage, ensuring only SLM-modulated components were transmitted. As a result, strict epifluorescence imaging as in Figure 2 A) i) is not used in our setup, instead small angles were employed to achieve similar effects. Lenses then relay light to the Olympus 60x NA 1.49 TIRF objective (Olympus APON60XOTIRF) such that both angled illumination and multifocal excitation were possible at the objective focal plane. All experiments in this paper used this objective. In both multifocal excitation and HILO and TIRF inclined excitation, emitted fluorescence is detected through same objective through a microscope frame (Olympus IX73P2F). A dichroic (Semrock) transmits the fluorescence light while blocking laser wavelengths. The tube lens then imaged the sample to the scientific CMOS camera (Andor), with additional notch filters used to block stray excitation light. The final arrangement of lenses leads to a demagnification from SLM to objective focal plane of 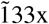 (one 9 *μ*m pixel on the SLM is 67.7 nm at objective focal plane). The full field of view of our system is 80×136 *μ*m.

We first tested the performance of the SLM for two lasers of 488 nm and 561 nm wavelengths. In the case of a flat phase profile which corresponds to *T* = 0 for *ϕ*_*blazed*_ described earlier, both light sources come to a common focus at the focus of L3. Increasing *T* produces a wavelength-dependent change in position of the focus. We characterised this position to relate the translated distance, D, to T via a linear equation of form *D* = *mT*. We derived *m*_488*nm*_=5.80 ± 0.004 *μm* / Δ*T* and *m*_561*nm*_ = 6.72 ±0.007 *μm* / Δ*T*, showing that 488 nm focus is translated a greater distance than the 561 nm focus. This matches expectation as a phase delay of 488 nm is equivalent to 2*π* radians for the 488 nm light, but only 488/ 561 × 2*π* radians for the green laser. This must be taken into account when designing phase masks at multiple wavelengths.

Secondly, we considered consequences of the spatial bandwidth of the SLM and aliasing. The SLM pixelation introduces a limit to the spatial frequencies in phase masks, which limits the achievable focus translation *D*. This can lead to generation of additional higher order intensities in the Fourier plane. In addition, the pure phase modulation of the SLM results in ghost orders, for which additional care must be taken [22–24]. There is also the presence of the zero order intensity which cannot be reduced below the pixel fill factor, leading to an on-axis focal position in all cases [25]. In our situation as in Figure 3 A), separate inspection cameras were used to characterise the above effects and SLM parameters were chosen to avoid unwanted intensities. In Supplementary Figure S5 we illustrate the importance of tailoring the light intensity in the BFP and how it affects HILO and TIRF image quality. TIRF is particularly sensitive to the presence of additional intensities as even a small component of non-evanescent excitation significantly reduces signal to noise and sectioning. We employed an indirect protocol to validate TIRF operation using a solution of sub-diffractive beads. The beads resting on the surface are brought into focus, while the SLM changes excitation inclination angle from epifluorescence and HILO to TIRF. In TIRF mode, the beads in solution are no longer excited and their movement cannot be seen apart from short moments when beads in solution hit the coverslip. Hence, regardless of theoretical prediction, BFP intensity must be checked for each SLM phase mask and appropriate spatial filtering and verification must be conducted.

## 3. Results

### 3.1. Characterisation of bleaching foci in uniform fluorescence solutions

We first studied at a uniform sample of 250 µg/mL fluorescein in 100% glycerol, to establish that both bleaching, and recovery were possible with the system. We applied dried 50 *μ*m beads to a coverslip before adding the glycerol solution. We then applied an additional coverslip to flatten and spread the glycerol layer. The beads ensured a separation of 50 *μ*m between the two coverslips, and were not directly used in the imaging process. Using 488 nm excitation, a single focus was generated, and its power modulated to switch between bleaching (high power) and recovery (low power). A focus at low power (0.5 mW) initiated the experiment and established the baseline intensity (I_initial_). Then, the spot was switched to the bleaching power (P_bleach_) for 10 seconds. Next, the spot switched back to low power and recovery was observed back to a steady state.

In Figure 4, we can observe the focal spot generated by the SLM for both Figure 4 A) i) photobleaching and Figure 4 A) ii) photorecovery. P_bleach_=5 mW in this example, which is sufficient to bleach the sample as seen when the fluorescent signal decreases from t1 to t3. In the recovery phase, we use low power (P_low_=0.5 mW) to visualise a recovery in the fluorescence signal. We see increasing intensity from t4 to t6. The image contrast has been adjusted for the high and low power mode examples, so it is not possible to compare the images in Figure 4 A) i) and Figure 4 A) ii) directly. A fixed line profile was selected across both the bleaching and recovery phases, and this was visualised over time in a kymograph (A iii). Each pixel along the horizontal axis represents the intensity in 10 ms increments, with the vertical axis showing the full line profile. The line profile along the time axis was then plotted to derive the full intensity curve in Figure 4 B).

**Fig. 4.**
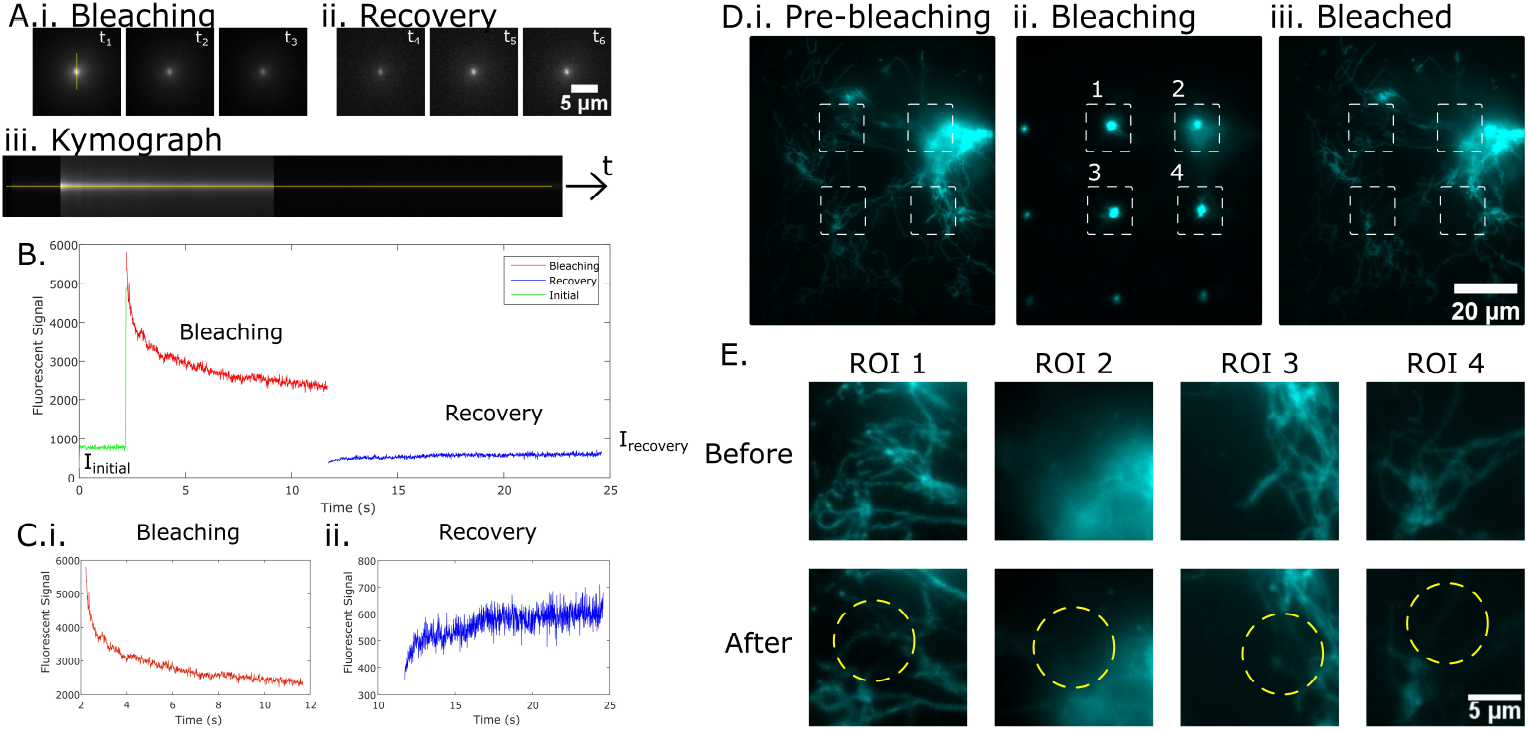
FRAP on a viscous fluorescent solution. A) i) Bleaching shown using a fluorescein-glycerol solution, and ii) subsequent recovery. iii) is a kymograph of the line profile. B) The full bleaching-recovery curve, with enlarged versions in C) i) and ii). D) sample of HMVEC endothelial cells stained with Mitotracker Orange, i) prior to photobleaching in HILO imaging, ii) during photobleaching, iii) After photobleaching. E) shows the zoomed in regions highlighted in the first images. ROI 1, 3, and 4 show very clear photobleaching go the mitochondrial structures (at yellow circles). ROI 2 shows that even mitochondria out of focus experience photobleaching, indicating the axial extent of the micro foci generated using the SLM.

### 3.2. Bleaching at multiple locations in extended biological samples

Figure 4 B) shows three distinct regions, comprising an initial steady state using low power (P_low_=0.5 mW), a bleaching phase with (P_bleach_=5 mW), and a recovery phase with (P_low_=0.5 mW). The bleaching and recovery curves in Figure 4 B) match the expected exponential decay and recovery. Given the large intensity difference between P_bleach_ and P_low_, we show each distinct phase in Figure 4 C) i) and ii). We then examined the bleaching and recovery rates for each of the curves. Three experiments each were conducted for P_bleach_=5 mW and P_bleach_=10 mW, with P_low_=0.5 mW for both cases. The bleaching rates were expected to be influenced by P_bleach_, but the recovery rates based on P_low_ was kept constant.

The bleaching and recovery curves were both fit to a two-parameter exponential fit as in Equation 3.

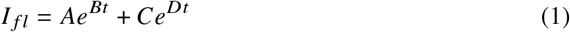

The parameters in the exponent B and D correspond to the rate of bleaching and recovery in the curves. Higher B and D values indicate a higher rate of bleaching or recovery. The parameters A and C are scaling constants that indicate the fluorescence involved.

We used the Matlab curve fitting toolbox to treat the bleaching and recovery curves separately for each experiment. A summary for the average rates for each experiment are presented in Table 1, and in text we refer to each line of the table with the convention (B, D).

**Table 1.**
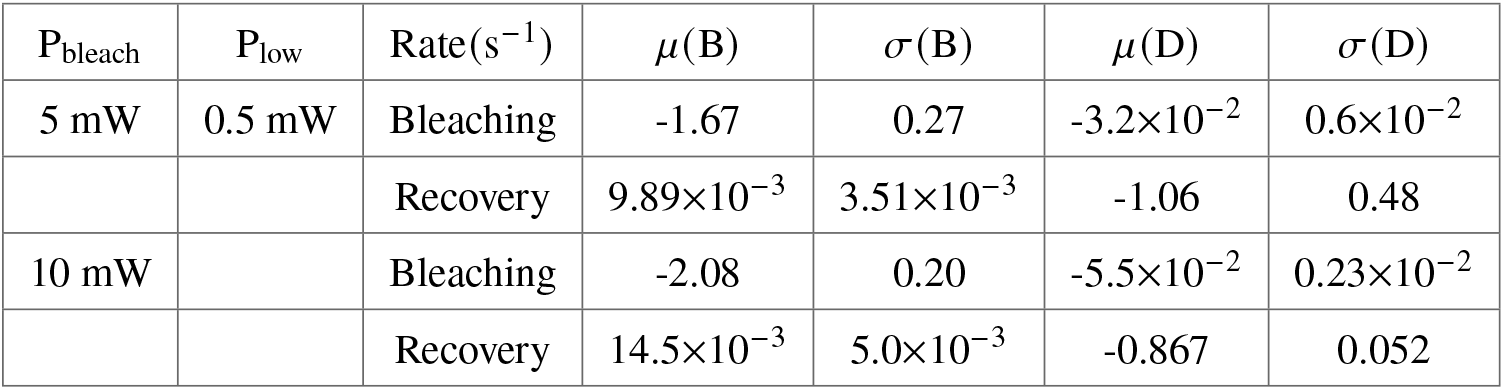
Experimental Results for Bleaching and Recovery.

| $P_{\text{bleach}}$ | $P_{\text{low}}$ | Rate($s^{-1}$) | $\mu(B)$ | $\sigma(B)$ | $\mu(D)$ | $\sigma(D)$ |
| --- | --- | --- | --- | --- | --- | --- |
| 5 mW | 0.5 mW | Bleaching | -1.67 | 0.27 | $-3.2 \times 10^{-2}$ | $0.6 \times 10^{-2}$ |
| | | Recovery | $9.89 \times 10^{-3}$ | $3.51 \times 10^{-3}$ | -1.06 | 0.48 |
| 10 mW | | Bleaching | -2.08 | 0.20 | $-5.5 \times 10^{-2}$ | $0.23 \times 10^{-2}$ |
| | | Recovery | $14.5 \times 10^{-3}$ | $5.0 \times 10^{-3}$ | -0.867 | 0.052 |

Under a 5 mW bleaching power, we measured a bleaching rate of (−1.67± 0.27, (3.2 ± 0.6)×10-2), while under 10 mW bleaching power, we measure (−2.08 ± 0.2, (−5.5 ± 0.23)×10-2). The higher bleaching power causes a more rapid photobleaching rate, and this is seen in higher B and D values. For the dataset using P_bleach_=5 mW, we obtain ((9.89 ± 3.51)×10-3, −1.06 ± 0.48) and ((14.5 5)×10-3, −0.867 ± 0.052) for the P_bleach_=10 mW dataset. Our measurements of bleaching and recovery rates are based on a simple two-term exponential model, and we show that this is sufficient to give a relative measurement of diffusion rate. The difference between I_initial_ and I_recovery_ values can be attributed to photobleaching leading to a smaller pool of fluorescent molecules.

Modelling solutions are available that account for boundary conditions and the intensity profile of the bleaching spot [26]. In this study we considered only Fresnel lens equations which effectively generate Gaussian foci. In principle it is possible to replace *ϕ*_fresnel_ to control the shape of each focus independently. For example, it is possible to incorporate aberration correction or other types of optical elements like micro axicons to control the bleaching profile (see examples in Supplementary Figure S4). Such control over the shape of each focus could lead to improved modelling for diffusion rate.

In this work we introduce a novel modality, which makes use of an SLM to dynamically switch between different widefield fluorescence imaging modes (epi, HILO, TIRF) and individually addressable multiple focal spots for photobleaching. The resulting instrument is uniquely suited to Fluorescence Recovery After Photobleaching (FRAP) studies. We expect this system to be beneficial for a range of protein diffusion studies, allowing the ability to distinguish between e.g. the cell membrane and the cell nucleus at camera-limited rates. Such a system combining such single molecule imaging capabilities (HILO) and FRAP can have strong utility as high speed switching between FRAP and HILO can overcome many of the limitations of scanning-mirror based FRAP. On top of that, the typical FRAP methods are limited to single points while this method can support multiple individual spots in parallel. For example, one could study multiple locations in a three-dimensional volume while using HILO to probe structures such as the nucleus further from the coverslip, and TIRF to image the cellular membrane with high contrast. Measuring multiple FRAP sites under HILO imaging will be especially useful to determine Arp2/3 complex-dependent actin networks that drive retrograde actin flow [27].

## 4 Author Contributions

Conceived project and research: AU and WML. Designed and implemented the optics and software for the HILO/FRAP system: AU. Performed characterization and demonstration experiments: AU and YJL. Generation and testing of phase masks: AU. Provided glycerol and cell samples and generated data for Figure 4: YJL. Data processing and instrument optimization: AU. Wrote paper: AU with input from all authors. Supervised research: WML.

## 5 Acknowledgements

We thank Hari Shroff (Janelia Research Campus, Howard Hughes Medical Institute) for discussions on the phase masks and comments on the paper. We acknowledge support from the Australian Research Council (DE160100843, DP190100039, and DP200100364), and NHMRC (APP2000485).

