## Supplementary Figures for "Combining Multi-site FRAP and HILO-TIRF microscopy using a Spatial Light Modulator"

### Simultaneous FRAP at multiple locations for high resolution kinetics measurement

This document provides supplementary information for the paper. Figures and text are derived from [1].

#### 1. MICROLENS FOCI IN OPTICAL TRAIN

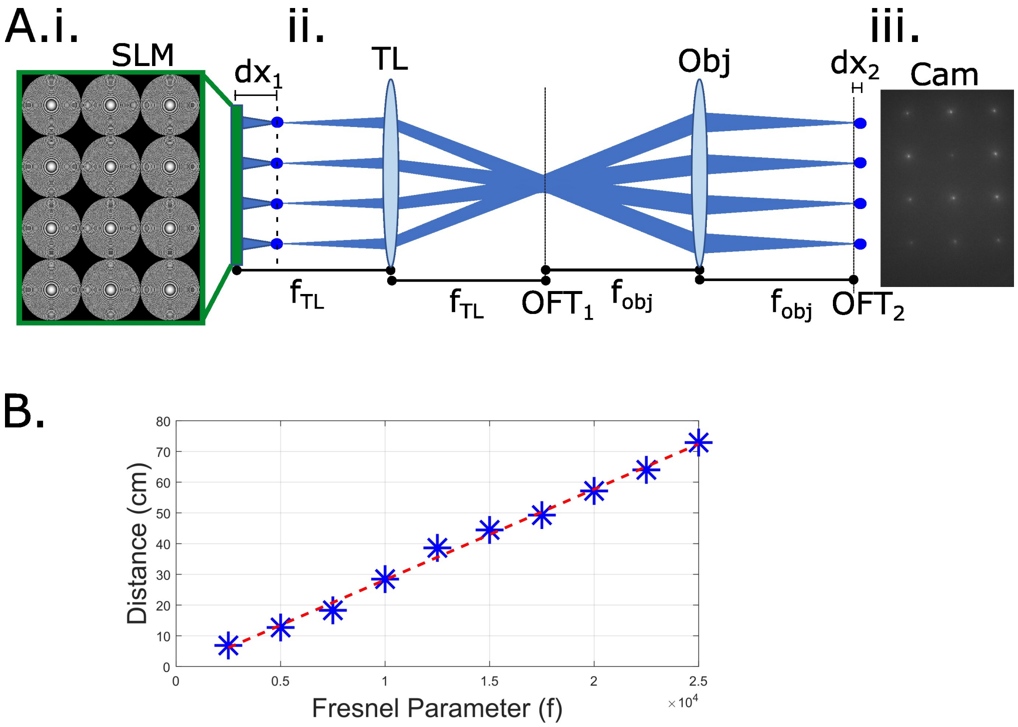


**Fig. S1.** SLM has individual control over foci. A) Fresnel lens mask on SLM produces array of microfoci at a distance dx from OFT2, achieved with 4f configuration. iii) intensity foci at sample. B) Changing the Fresnel parameter (f) gives linear change in focal position dx1, which is linearly mapped to dx2.

#### 2. SLM HAS INDIVIDUAL CONTROL OVER FOCI

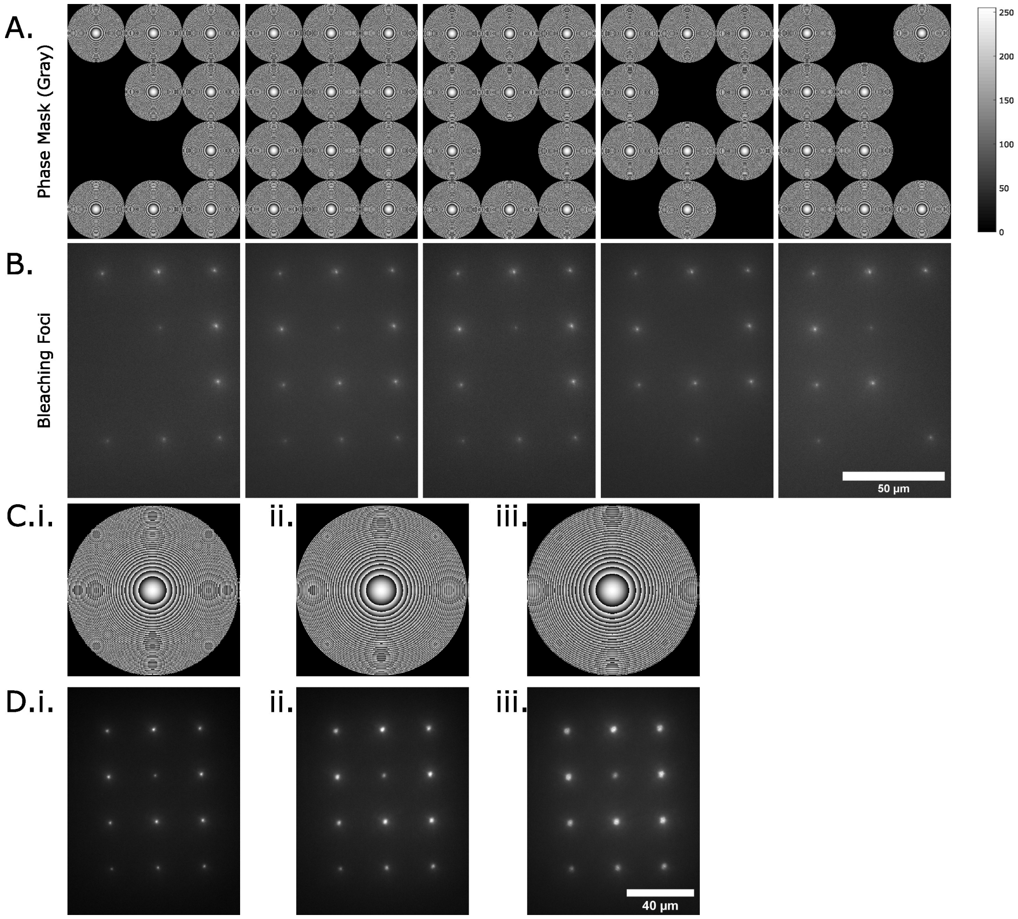


**Fig. S2.** Demonstrating individual control over the foci.In A) we see that the micro fresnel phase masks can be switched on and off at the SLM to determine whether a spot forms at the OP. In C) we incrase the Fresnel focal parameter on the SLM (one lens in the array shown) to produce D) successively defocused and broader focal profiles.

#### 3. SOFTWARE INTERFACE

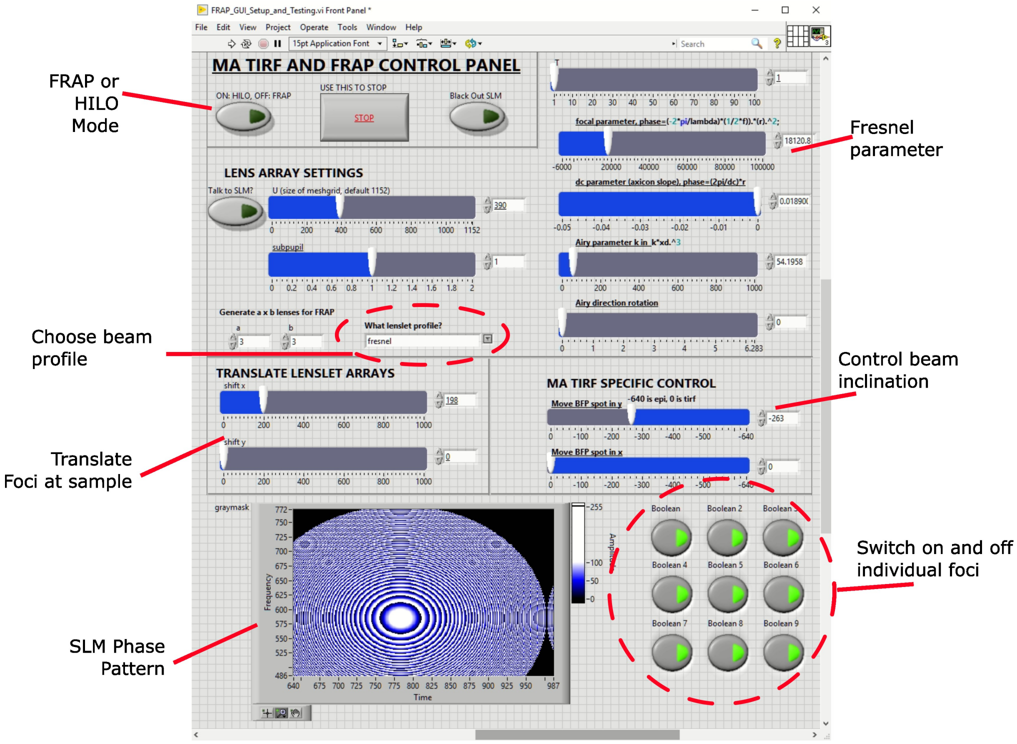


**Fig. S3.** The user interface for the multi-FRAP/HILO system. The system is fairly simple to use after initial setup. For FRAP, the system can control beam profile, focal length (axial position of foci at sample), switching on and off foci individually, and position of the arrays. For widefield imaging mode, the system allows switching between HILO and TIRF modes.

#### 4. BEAM PROFILES GENERATED BY SLM PHASE MASK

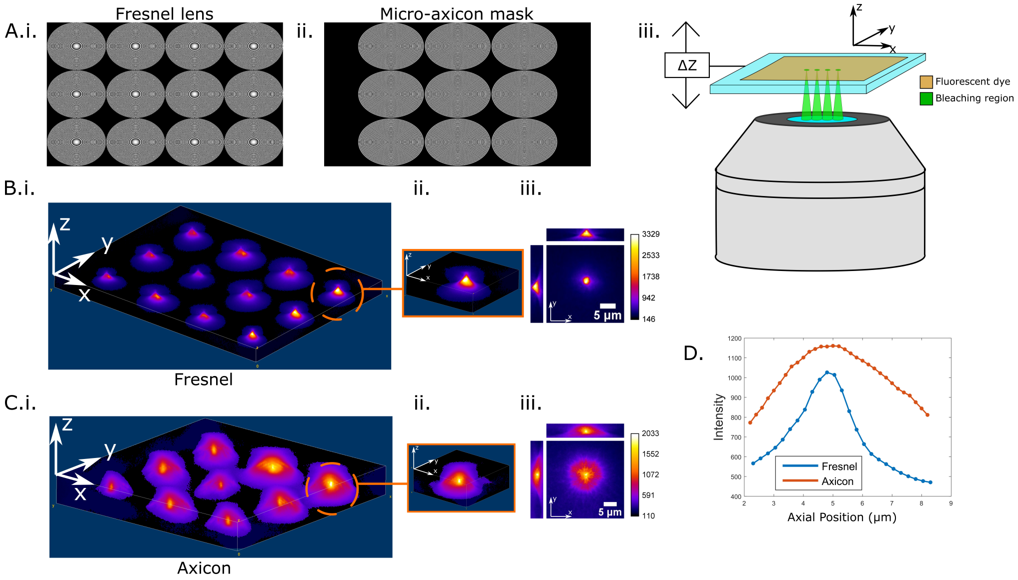


**Fig. S4.** Modulating the beam profiles. A) SLM phase masks for i) Fresnel lenslets and ii) Axicon profiles. The fluorescent intensity profiles at the sample plane using a single layer fluorescent sample for B) Fresnel lenslets and C) Axicon profile. In both cases we can see that there is a peak in intensity with different signal to noise ratios.

#### 5. TIRF AND HILO QUALITY

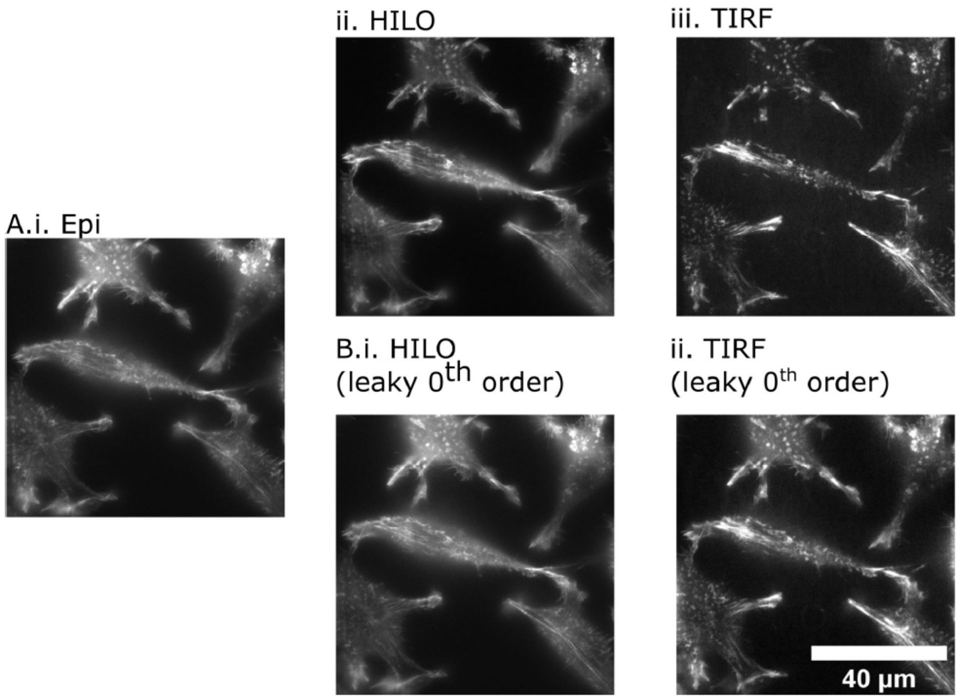


**Fig. S5.** Widefield fluorescence imaging using SLM. L929 mouse fibroblast cells fixed and labelled for F-actin (phalloidin). A) i) Epifluorescence, ii) HILO, iii) TIRF for same FOV. B) The presence of additional intensities in the BFP (in this case zero order is not blocked) can lead to fringing effects in the excitation. i) Compared to A) ii), there is more background fluorescence and the image looks more like epifluorescence. In addition, coherent interference creates fringes as seen in the cell in the bottom right. B) ii) The zero order significantly reduces the signal to noise of TIRF as fluorescence deeper within the cell is excited. In this case, fringing effects are not usually seen.

In Figure S5 B) i), we see that not blocking the zero order will reduce SNR, as well as leading to fringing effects. This is a result of the zero order and the first order light from the SLM interfering coherently, with resulting fringes clearly seen in the bottom right cell. In Supplementary Figure S5 B) ii) the zero order is again unblocked but in the TIRF case. In TIRF, no beam should emerge from the objective as the light undergoes total internal reflection. Hence this modality is particularly sensitive to additional intensities. Any component that is not in TIRF mode, i.e. Epi-fluorescence, will significantly contribute to the background fluorescence. This leaking intensity removes the inherent advantage of TIRF – the axial sectioning. As expected the leaky TIRF image in Figure S5 B) ii) shows significantly lower SNR. These effects can occur not only from improper blocking of the zero order light, but aliasing effects described earlier.

When using blazed gratings, especially those with high period as in HILO and TIRF, the phase mask must be inspected to ensure there are no stray orders. This can be done through a camera in the BFP. Alternatively, for a visual check of the emerging oblique beams, a petri dish with water should be prepared. A highlighter can be dipped into the water, generating a fluorescent solution that allows for observation of the beam as it passes through. The illumination angle can be changed by generating the appropriate phase masks on the SLM. This will tilt the collimated beam, thereby changing the presence of evanescence/evanescent depth penetration. As the beam is tilted further, eventually no/minimal beams will be seen in the fluorescent dish – this indicates that HILO has ceased, and that Total Internal Reflection has been reached.

#### 6. HILO-FRAP ON LIVE CELL EXPERIMENTS

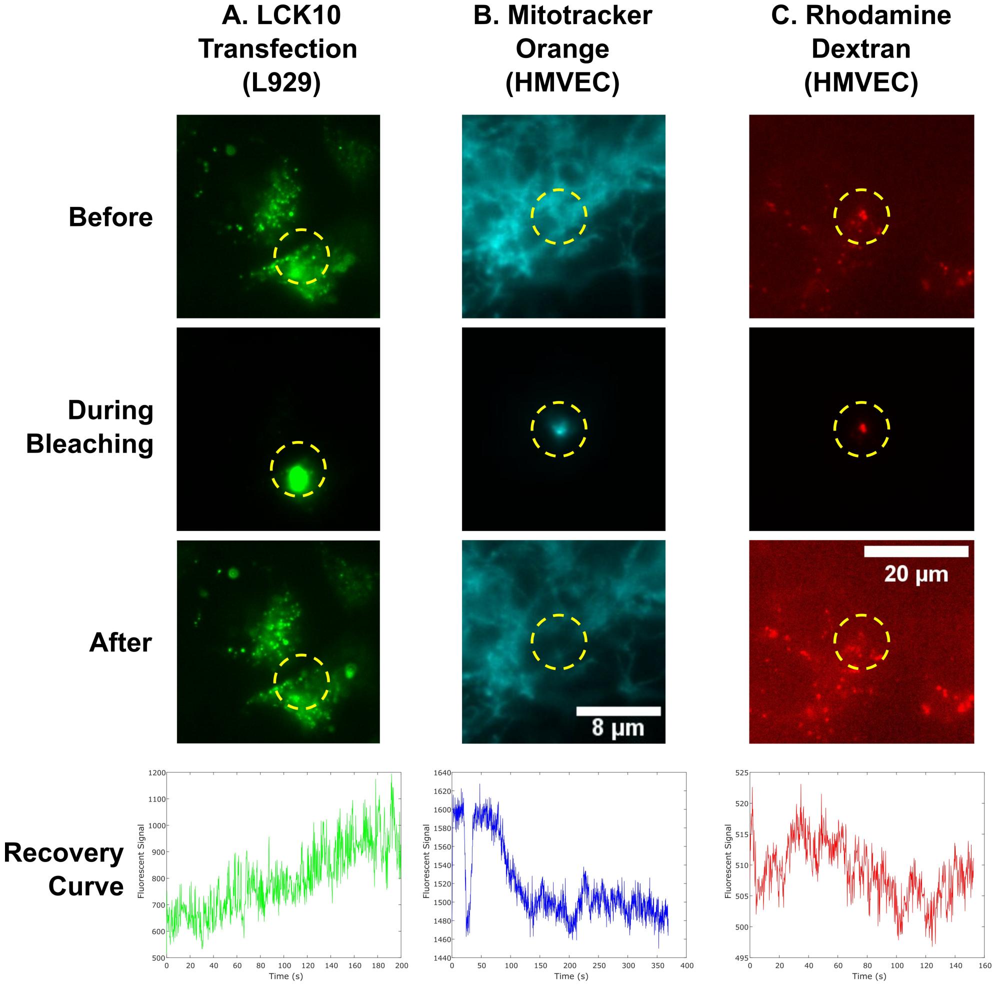


**Fig. S6.** Comparing three different fluorescent cell samples for A) L929 cells transfected with LCK10, a membrane protein. B) HMVEC cells stained with Mitotracker orange. C) HMVEC cells incubated with Rhodamine Dextran, a large polysaccharide (70 kDa). For each of these, we show photobleaching and the fluorescence curve after bleaching. We applied a moving average with window of 15 in each case to reduce the fluctuation of the signal.

We considered three separate fluorescent samples to illustrate the use of our HILO-FRAP system for live cells. Live cell samples are much more heterogeneous than the previous uniform glycerol sample. The complex intracellular environment scatters light, meaning we cannot assume photobleaching will occur as in the glycerol sample. In addition, the type of staining will affect the type of fluorescent intensities observed, and the degree to which their recoveries are influenced by diffusion. The goal was to determine whether FRAP measurements could be conducted on live cells, and whether the HILO system produced images with sufficient detail to visualise cellular environments.

We considered HMVEC endothelial cells stained with Mitotracker Orange, which is a mitochondria-specific stain that can be excited by our 561 nm laser. A 37° heat plate was used to maintain the cells for imaging, and the cells were visualised in glass bottom dishes (Cellvis). In Figure S6 D), we show HILO images of our mitochondria stained HMVEC cells with a 3×3 array of foci across the FOV. The mitochondria appear as long tube-like extensions that form a network throughout the cell. It is difficult to see the foci to the left and bottom of the FOV since they are not aligned to any fluorescent structures, so we focus on four central spots labelled 1-4. We show the FOV for Figure S6 D) i) before, ii) during, and iii) just after photobleaching, and present enlarged images in

Figure S6 E). In Figure S6 E), bleaching has occurred in four different locations (circled in yellow).

In ROI 1, the mitochondria intensity just after bleaching is clearly lowered in the centre of the image. This drop in signal in the centre compared to the outer edges shows the presence of targeted point photobleaching. There is, however, also a global drop in fluorescence which can be attributed to the long term HILO imaging. In ROI 2, we see a dense mass of out of focus structures. Bleaching in the centre of the image does occur, which indicates that our bleaching is not restricted to one plane. This is expected based on the focal profiling of the technique. In ROI3, we see that the outer regions of the ROI remain fluorescent, while the protrusion in the centre is photobleached. Finally, point bleaching is seen in ROI 4 again. These results indicate that the HILO-FRAP system can be practically used to visualise the intracellular environment of a cell, and conduct spatially selective photobleaching.
